# The Physical Effects of Learning

**DOI:** 10.1101/2023.06.23.546243

**Authors:** Menachem Stern, Andrea J. Liu, Vijay Balasubramanian

**Affiliations:** Department of Physics and Astronomy, University of Pennsylvania, Philadelphia, PA 19104; Center for Computational Biology, Flatiron Institute, Simons Foundation, New York, NY 10010, USA; Santa Fe Institute, 1399 Hyde Park Road, Santa Fe, NM 87501, USA; Theoretische Natuurkunde, Vrije Universiteit Brussel, Pleinlaan 2, B-1050 Brussels, Belgium

## Abstract

Interacting many-body physical systems ranging from neural networks in the brain to folding proteins to self-modifying electrical circuits can learn to perform specific tasks. This learning, both in nature and in engineered systems, can occur through evolutionary selection or through dynamical rules that drive active learning from experience. Here, we show that learning leaves architectural imprints on the Hessian of a physical system. Compared to a generic organization of the system components, (a) the effective physical dimension of the response to inputs (the participation ratio of low-eigenvalue modes) decreases, (b) the response of physical degrees of freedom to random perturbations (or system “susceptibility”) increases, and (c) the low-eigenvalue eigenvectors of the Hessian align with the task. Overall, these effects suggest a method for discovering the task that a physical network may have been trained for.

## I. INTRODUCTION

Nature is replete with systems that learn. For example, animals learn new behaviors, the immune systems of vertebrates and bacteria learn pathogenic environments, and, over evolutionary time, proteins learn structures that fold just right to achieve precise and specific molecular functions. In supervised computational machine learning [1, 2], the learning process is formulated as minimization of a *cost function* on a system’s input-output behavior. More generally, this formulation provides a powerful paradigm for solving difficult inverse problems [3, 4]. The same paradigm has also been exploited to describe biological learning in various forms, e.g., in neural [5] and immune systems [6].

Biological systems are necessarily physical in nature and therefore, like any physical system, must obey certain constraints that cause their learning processes to differ from those of computers. In particular, while computer algorithms often seek to globally descend cost function gradients with respect to learning degrees of freedom (e.g., neural network weights), physical systems without external processors cannot generally implement such optimization processes, even though learning by natural selection can be sometimes be cast in this way. Typical biological learning, e.g., by an animal learning a new behavior, operates on time scales far shorter than evolutionary ones, and must proceed by dynamical processes (learning rules) that modify internal (learning) degrees of freedom in response to examples [7]. Such learning rules are generally local in space and time and cannot be informed about the functionality of the whole system. In other words, learning in physical systems on shorterthan-evolutionary time scales differs from computational machine learning in that the learning is *emergent*. It is a collective behavior of many elementary units, each implementing simple rules based on its own local environment. The Hebb rule in neuroscience (“neurons that fire together, wire together”) is an example of a local rule – synaptic plasticity is based on local information, leading to debates about whether and how such dynamics propagate information about the training task to individual neurons and synapses. Local rules have also been exploited to train laboratory non-biological mechanical networks to exhibit auxetic behavior [8, 9] or protein-inspired functions [10–12]. Other local rules have been proposed for associative memory [13–17]; one of them has even been demonstrated in the lab [18]. Here we will focus on a powerful set of local rules based on the framework of Contrastive Hebbian Learning, which perform approximate gradient descent of a cost function [19–26]. Such contrastive learning was recently realized experimentally for tasks including regression and classification in electronic resistor networks [27, 28].

Here, we focus on physical systems such as athermal mechanical, flow or electrical networks, in which the learning rate is slow compared to the rate of physical relaxation. In this limit, mechanical networks remain in equilibrium during the learning process, while flow/electrical networks remain in steady state. As a result, forces on nodes of a mechanical network must add to zero, while all the currents through nodes of a flow or electrical network must add to zero. These constraints arise because the system must typically be at a minimum of a *physical* cost function (e.g., the energy in a mechanical network or the dissipated power in a flow/electrical network) with respect to the physical degrees of freedom (e.g., node positions in mechanical networks, or currents through edges of flow/electrical networks). These physical degrees of freedom couple the learning degrees of freedom (e.g., spring constants in mechanical networks or conductances for each edge in flow/electrical networks). Note that systems that operate at a minimum of a physical cost function are naturally recurrent in the sense that information flows in all directions, not only from ‘inputs’ to ‘outputs.’ Thus, they differ fundamentally from computational learning algorithms based on optimizing a feed-forward functional map.

In such equilibrium or steady-state systems, learning involves a *double* gradient descent – at every step of minimization of the learning cost function, the physical cost function must also be minimized with respect to the physical degrees of freedom. Therefore there are two landscapes of interest: the learning cost function in the high-dimensional space spanned by the learning degrees of freedom, and the physical cost function in the high-dimensional space spanned by the physical degrees of freedom. These two landscapes are coupled together. As a system learns, either directly by double gradient descent or by using local rules that approximate gradient descent in the learning landscape, changes in the learning degrees of freedom sculpt the physical landscape. We examine the effects of learning on this physical landscape and the properties of the learning system.

We are inspired by recent efforts in the fields of protein allostery and computational neuroscience that suggest learning has a set of typical effects on the physical structure of a network and its response to external forces. Some folded proteins, often modeled as mechanical networks [29, 30], have evolved allosteric function, where binding of a regulatory molecule at a “source” site triggers a conformational change in the protein that either enables or inhibits binding of another molecule at a distant “target” site. Model networks, as well as networks derived from folded proteins, that display allosteric behavior have been shown to exhibit low-dimensional responses to strains applied at the source [31–40]. In particular, the allosteric response is well-captured by low-energy (soft) normal modes of vibration [36, 37, 41, 42]. Similar observations have been made in neural circuits and model systems [43–47], in particular for unsupervised learning tasks such as predictive coding [48]. Our goal is to understand how such effects arise.

In this paper, we study the effects of learning on the physical landscapes of mechanical and flow networks, as they use local rules to learn tasks by effectively performing double gradient descent in the physical and learning landscapes. We show in detail how the structure of the physical landscape changes to accommodate the learned tasks, how it develops features observed in protein allostery studies [36, 37, 41, 42], and how it is affected by loading multiple concurrent tasks on the network, up to and beyond its capacity [49]. We find that, generically, when physical networks learn tasks in the linear response regime, they become soft as learning proceeds, and the responses become low-dimensional, aligning with directions of low curvature in the physical landscape. The tasks that can be learned in this way include not only allostery, as in proteins, but typical computer science tasks such as regression and classification. In a sense, we are showing that *things become what they learn*. The implication is that learning imprints signatures on a physical system, allowing an external observer to gain insight into whether a network has been trained, and for what tasks.

## II. TRAINING A PHYSICAL NETWORK

Consider a network (Fig. 1a) with physical degrees of freedom *x*_*a*_, *a* = 1 … *N* collected into a vector 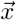, and learning degrees of freedom *w*_*i*_, *i* = 1 … *N*_*w*_ collected into a vector 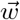. We apply inputs to one subset of the physical degrees of freedom, and designate another subset as outputs. Learning modifies the learning degrees of freedom to improve the physical responses of the outputs, driving them closer to the desired responses. For an athermal mechanical spring network, we will designate the positions of the nodes as physical degrees of freedom {*x*_*a*_} that adjust to minimize the elastic energy, which is the physical cost function *E*. The learning degrees of freedom {*w*_*i*_} are spring stiffnesses [10, 49]. The inputs are forces on nodes and outputs are node displacements. For a flow network, {*x*_*a*_} represent the node pressures, which adjust to minimize the power dissipated, which is the physical cost function *E*. The learning degrees of freedom {*w*_*i*_} are the conductances of the edges of the network while the inputs are externally applied currents at the nodes [49] and outputs are node pressures. In an electric resistor network, {*x*_*a*_} may correspond to the node voltage values while {*w*_*i*_} correspond to the edge resistances [23, 27].

**FIG. 1.**
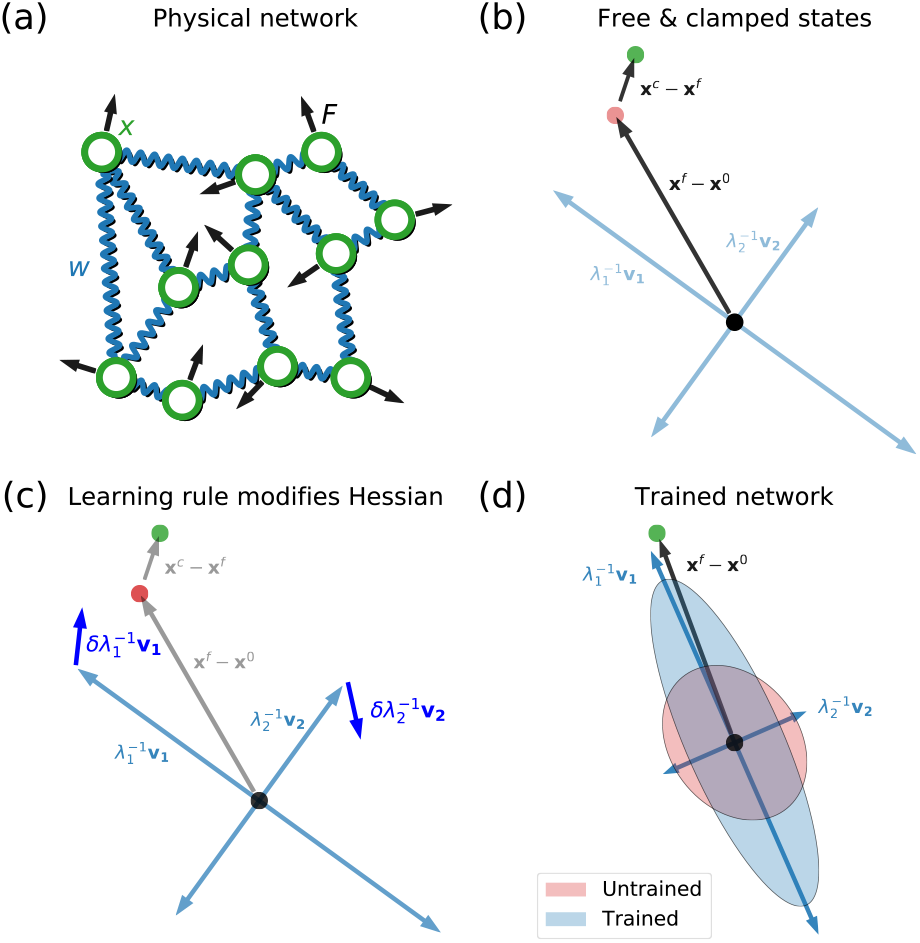
Learning modifies the physical network. a) Input forces (black) are applied to the physical degrees of freedom (green, e.g. node positions) of a physical network (e.g. mechanical spring network), whose interactions correspond to learning degrees of freedom (blue, e.g. spring constants). b) In the physical configuration space, this input force causes the system to respond, equilibrating in a *free state* (red dot). To train the system, a further ‘output force’ is applied, nudging the system to a *clamped state* (green dot). The blue arrows describe the inherent physical coordinate system, with direction corresponding to eigenmodes *v*_*ab*_ of the physical Hessian *H*_*ab*_, and lengths correspond to the associated inverse eigenvalues 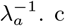) A local learning rule is applied, modifying the learning degrees of freedom. On top of improving the system free state response, learning tends to rotate the Hessian coordinate system such that the eigenmode corresponding to the lower eigenvalues align with the free state response, and decrease these eigenvalues. d) Training results in a physical system whose lower eigenvalues are reduced, and eigenmodes aligned with the trained task(s). The system responds considerably more strongly to random forces, shown by the area spanned by the trained inverse eigenvalues (blue ellipse) compared to the untrained ones (red ellipse). Training makes the physical system more conductive and lower dimensional.

Protocols for deriving physically realizable learning rules for such systems can be constructed based on the ideas of Contrastive Hebbian Learning [19]. These approaches include Equilibrium Propagation [20], Coupled Learning [21] and Hamiltonian Echo Backpropagation [22]. All of these local learning rules are based on the comparison of two states of the system: (a) a *free state* where only inputs are applied, and (b) a *clamped state* where the outputs are nudged toward the desired values (Fig. 1b). Below, we show how such learning rules change the physical system (Fig. 1c), aligning its response eigenspace with the learned task, and increasing its responses to random forces (Fig. 1d).

More precisely, consider a noiseless physical network with fixed learning degrees of freedom 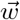. The network dynamics and responses to external inputs are fully determined by the physical cost function 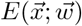 which controls how the physical degrees of freedom 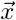 respond to external inputs. In the absence of such inputs the system equilibrates, locally minimizing the physical cost function to settle into a *native state* 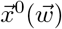 in which forces (for mechanical networks) or currents (for flow or electrical networks) are balanced 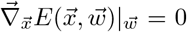, or, in components, 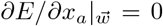 for all *a*. Below we will consider small perturbations around such native states; that is, we discuss learning in the limit of linear response. In fact, such learning approaches are practical and successful well beyond linearity [21]. However, in the linear regime we will show that learning is dominated by characteristic network phenomenology including reduction of the effective dimension of responses, softening of the system, and alignment of dynamics with the learned task. Beyond linear response, other mechanisms, such as multi-state learning [16, 42, 50], also become relevant.

In linear response it suffices to consider an expansion of the physical cost function around the native state up to the first non-vanishing, i.e., second, order:

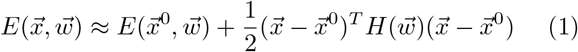

where the superscript *T* denotes the transpose, and *H* is a (symmetric) *physical Hessian* matrix, the components of which are 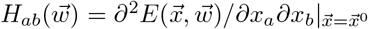. In the following we name this matrix the Hessian as a shorthand. The first order term in the Taylor expansion vanishes, since the native state 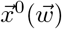 is a minimum of the physical cost function. The Hessian is a function of 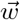 both explicitly, and implicitly through the dependence of 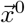 on the learning degrees of freedom 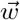.

To simplify the language in the remainder of this paper, we will use “force” to denote forces in the case of a mechanical network or currents in the case of a flow network.

### A. Training physical responses at network nodes

Consider a generalized external input force 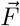 applied to the physical degrees of freedom, namely tensile forces in a mechanical network, or a set of currents in a flow network, applied to specific nodes. This force could be applied locally (at a subset of nodes) or globally, e.g., a “compression” applying forces to all the 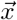 toward a certain point. These input forces will affect the physical cost function, prompting the system to equilibrate in a new *free state* 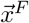 which minimizes the free state physical cost function:

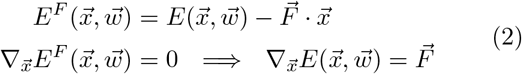

In the linearized approximation (1), this gives

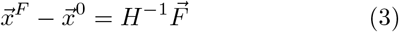

where we used the fact that the Hessian matrix is symmetric. In other words, the system responds by shifting the native state by the inverse Hessian applied to the input force. Note that 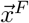 depends on the learning degrees of freedom 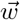 explicitly through the Hessian, and implicitly through the native state 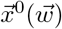.

Thus, deformations around the native state with small Hessian eigenvalues are “soft” – they exhibit a larger response to applied forces. Note also that the entries of the inverse Hessian depend globally on all the learning parameters. So the change produced by an external force on a given physical degree of freedom can depend nonlocally on the values of all the learning degrees of freedom, and not just on, say, the weights of edges connected to the network node in question.

In order to proceed, we must define a task in terms of desired outputs in response to the inputs, which we can express in terms of physical constraints that must be satisfied. The constraints can be local, applying to a subset of nodes designated as output nodes, or global, applying to all of the nodes. Because the response is expressed in terms of the physical degrees of freedom, the constraints can be defined by demanding that the free state response 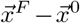 satisfies a desired relationship encoded in a functional 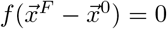. For example, if one desires 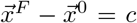 at a given output node *o*, then the functional is simply 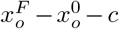. This can be thought of as a single basic task. More complicated tasks can be defined by adding functionals for many such tasks. For example, a classification task may be posed such that all members of each class (each set of inputs in the class) prompt a response that satisfies a particular constraint for that class.

Suppose we have multiple tasks, i.e., pairs of input forces and output constraints: 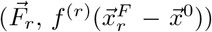 indexed by *r* = 1 … *n*_*T*_. We can quantify how well the system performs the tasks, i.e., implements these response constraints, in terms of a *learning cost function C*. For example, we can use a Mean Squared Error (MSE) cost:

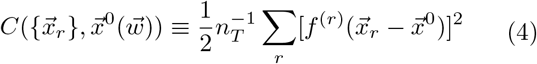

evaluated at the free state responses 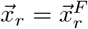. For a random, untrained physical system, the free state responses are completely independent from the desired responses, so the learning cost function *C* will tend to be high. Learning is the process of modifying the system, changing its learning degrees of freedom such that the cost function is reduced. How can this be done?

Computational machine learning algorithms generally minimize the learning cost function by performing gradient descent on *C*. Typically, this means that global information is required to determine local changes in the network. By contrast, a physical learning system must use local learning rules to achieve the same goal. We take the approach of Contrastive Learning in its incarnation as Equilibrium Propagation [20].

First consider training a system for a single task defined by the target constraint 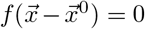. Suppose we apply the input force 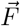 and the system equilibrates at some free state 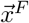. To train the system we could nudge it towards the desired output by applying an additional weak *output force*

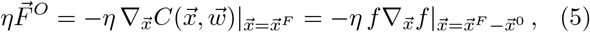

where *η* is a small parameter that we have explicitly separated out, and the last equation applies to a quadratic cost function (Eq. 4). The linearized system responds to the nudge by settling into a *clamped state* 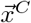 satisfying

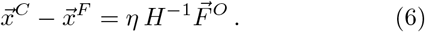

The clamped state depends on the learning degrees of freedom 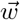 explicitly through the Hessian and implicitly through the additional dependencies in the free state 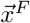. We can describe this equivalently by saying that the system minimizes a *clamped* physical cost function

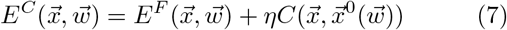

Thus the clamped state is the free state, nudged slightly by an extra output force related to a learning cost that arises if the system does not satisfy the desired constraints.

The contrastive learning [19] approach, later refined by equilibrium propagation [20] and coupled learning [21], compares the free and clamped states to derive an approximation to the gradient of the learning cost function that can be minimized more readily via local learning rules. Define the contrastive function:

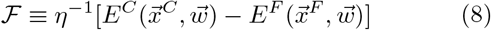

Previous work has showed that the partial derivative of the contrastive function with respect to the learning degrees of freedom 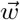 approximates the gradient of the learning cost function *C* in the limit *η* → 0 [20]:

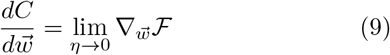

On the right hand side we differentiate only the explicit 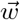 dependencies in the contrastive function, and not the implicit dependencies via the solutions for the free and clamped states. We will also assume that the MSE cost function *C* does not depend directly on 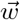, as it is a combination of physical constraints. The only explicit dependence of the physical cost function on the learning degrees of freedom then appears in the physical Hessian 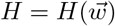.

Using (2) and (7) for *E*^*F*^ and *E*^*C*^ in ℱ, the Taylor expansion of the physical cost function in (1), and the linearized approximations for *x*^*F*^ and *x*^*C*^ in (3) and (6) gives

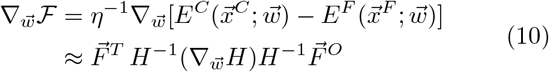

where we kept only the term that survives the *η* →0 limit in the second line, and used the fact that *H* and *H*^*−*1^ are symmetric matrices.

Learning now proceeds by following this derivative of the contrastive function with a learning rate *α*

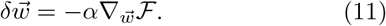

This learning rule has two key properties. First, learning is local. Every learning degree of freedom is modified according to the local difference between the free and clamped values of the physical cost, spatially localized at that edge of the network [20, 21]. This locality is a direct consequence of the property ℱ≥ 0 so that it does not have to be squared as in the learning cost function *C* in order to guarantee that the global minimum is at ℱ = 0. Second, the rule for modifying 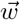 is proportional to the alignment of the input force 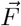 and the output force 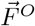 in an inner product determined by the symmetric matrix 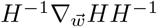. Each learning degree of freedom tends to decrease if the input and output forces align, and increase otherwise. We will see later how this property causes a realignment of the inherent physical coordinate system.

To illustrate, consider a model where 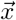 are physical variables at *N* nodes, while the 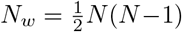 learning parameters are at edges linking each pair of nodes. In this fully connected case, the learning parameters are naturally represented as a symmetric matrix *W* with entries *W*_*ab*_, where *a, b* = 1 … *N* (Fig. 2a). The energy (physical cost) function for such a *node network* is a function of the node variables:

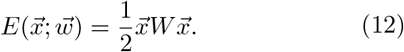

The Hessian of the energy is *H* = *W*, and is thus linear in the learning degrees of freedom. For sufficiently small forces compared to the scale set by the Hessian, any classical Hamiltonian system can be written in this way, and learning is the process of modifying the edge weights to amplify desired responses around rest, while suppressing unwanted responses to inputs. For this model the learning rule for a given edge weight *W*_*ab*_ is

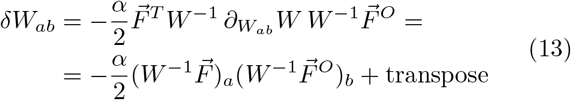

Here, we used the fact that the *a, b* and *b, a* entries of the matrix 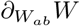 equal 1, while all the other entries are 0. So in the second line we are multiplying the *a*^th^ component of the vector 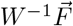 by the *b*^th^ component of 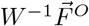 and adding the transpose.

To show that the physically realizable learning rule Eq. 13 can converge, we first tested it on a relatively easy task. We initialized networks with *N* = 20 nodes with energy functions of the form (12) and weights *W*_*ij*_ drawn randomly from a standard Gaussian 𝒩(0, 1). To ensure a positive semi-definite Hessian, we explicitly symmetrized the weights and added a term proportional to the identity 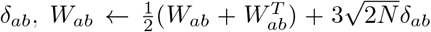. Then *H* = *W* (since W is already symmetric) and all its eigenvalues are positive. The resulting random Hessians initially have eigenvalues in the range ∼ (10, 27). We then defined a random learning task by picking an input force with each component drawn from a standard Gaussian, and a linear output constraint 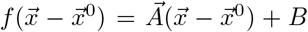 with the entries of 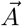 and *B* also drawn from standard Gaussians (see Appendix A). The local learning rule was effective in training such networks, reducing the error in Eq. 4 by many orders of magnitude (Fig. 2b). Appendix A gives examples of other tasks such as allostery, regression and classification that can be trained in this way.

**FIG. 2.**
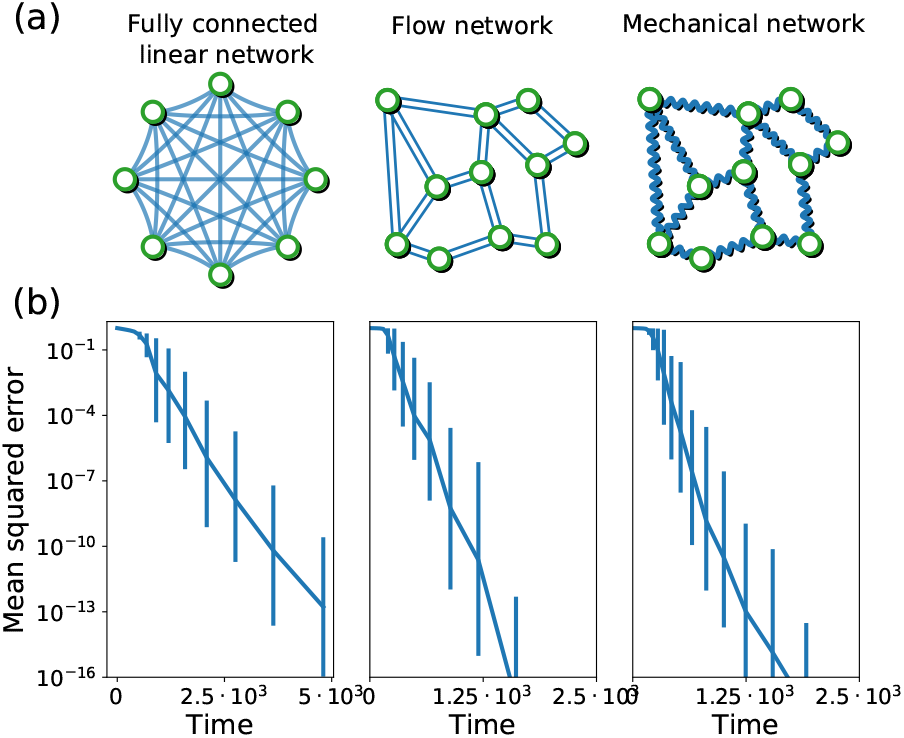
Training physical networks. a) We train several types of physical networks in the linear response regime, in particular fully connected linear networks (where physical degrees of freedom are node values), Flow networks and mechanical spring networks (where physical responses are naturally represented as difference over edges). b) Training these systems on single tasks, we generally find that local learning rules succeed, reducing the mean squared error by many orders of magnitude (geometric mean over 50 realizations of networks trained with single tasks).

### B. Training mechanical and resistor networks

In the mechanical/flow/electrical-resistor networks of interest to us, the physical cost function minimized by the network is defined via differences in the physical degrees of freedom 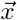 between nodes connected by edges whose non-negative learning degrees of freedom are 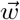. Each element *x*_*a*_ of 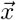 can itself be an element of some d-dimensional vector space describing the physical attributes of nodes. For example, we will consider mechanical networks where the node variables *x*_*a*_ correspond to positions 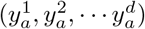 in a *d*-dimensional physical space, while the weights *w*_*i*_ are spring constants (stiffnesses). We will use the notation that the product of two node variables *x*_*a*_*x*_*b*_ should be understood as an inner product in the physical space. For example, if the physical space is d-dimensional Euclidean space then 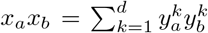. Likewise in resistor/flow networks, the *x*_*a*_ describe node voltages/pressures (*V*_*a*_), which we will regard as one-dimensional vectors, and the *w*_*i*_ describe conductances on edges. The energy (power) in these networks is a weighted sum over edges of the squared strains for springs, or squared voltage drops for resistors. In view of this, it is more convenient to work with the *differences* between physical node variables connected by edges (Fig. 2a) rather than the node values themselves.

To this end, we arbitrarily assign an orientation to every edge and define Δ_*ia*_ to take the values ±1 depending on whether the node *a* is at the incoming or outgoing end of edge *i*, and 0 otherwise. The *N*_*w*_ × *N* quantities Δ_*ia*_ form the *incidence matrix* Δ, such that 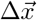 is an *N*_*w*_ dimensional vector, with entries that are the differences between the node variables on either side of each edge. In terms of this incidence matrix, we can write the physical cost function minimized by such *difference networks* as

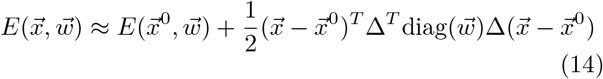

where 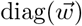 is a diagonal matrix with entries *w*_*i*_ which measure the conductances (for resistance/flow networks) or stiffness (for central-force spring networks) of the *N*_*w*_ edges. Here, if the edge *i* is incident on nodes *a* and *b*, then the *i*^th^ component of 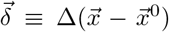 is 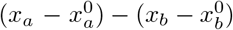. In terms of these differences the second term in the physical cost function (14) is ∑_*i*_ *w*_*i*_*δ*_*i*_ *· δ*_*i*_. Since the *δ*_*i*_ are differences of d-dimensional physical node variables, as described above, their product is defined by an inner product in the physical space. We can map the difference network (14) to the node network (12) by the identification

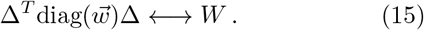

We can thus identify the physical Hessian as 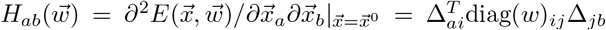 with 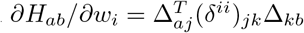 where *δ*^*ii*^ is a *N*_*w*_ × *N*_*w*_matrix with a 1 in the *i*^th^ diagonal entry, and zeros elsewhere. We similarly define the learning task as the satisfaction of a linear constraint, 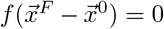. In terms of these constraints we define a learning cost function *C* = 0.5*f* ^2^. Similarly to Eq. 3, given an input 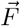, the free state response is 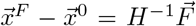. The clamped state response is also the same as before, 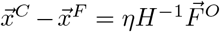, with an output effective force 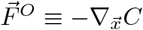. In mechanical spring networks, such forces cause the displacement of network nodes, while in flow or resistor nets, these forces may be understood as injecting currents to nodes. Using these definitions and Eq. 11, we find that the learning degrees of freedom 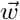 change in a training step by

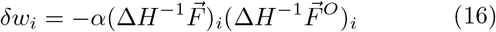

where on the right side we are simply multiplying the *i*^th^ component of the vectors 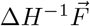 and 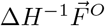. Overall, learning in such physical networks where the relevant degrees of freedom are differences of node variables is similar to the networks discussed above in which the physical variables are the node variables themselves. In particular, (16) could be rendered in matrix form like (13) if desired, except that the entries of the *W* matrix in this case would be linear combinations of the network edge weights dictated by the incidence matrix. We also see that *δw*_*i*_ in (16) can be written as minus the product of the *i*^th^ components of the free and clamped state displacement difference at the *i*^th^ edge. This means that the conductance/stiffness elements 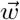 tend to decrease if the response to the input and output forces at an edge align and increase otherwise.

In Fig. 2b, we show that Eq. 16 successfully trains *N* = 40 flow networks and elastic spring networks for allosteric tasks (see appendix A), reducing their error by multiple orders of magnitude. These networks were derived from Erdőos–Rényi graphs with mean coordination number *Z* = 3 for flow networks and *Z* = 4 for 2-dimensional mechanical networks.

In summary, in the remainder of the paper we will focus on these three example types of networks. (1) A fully-connected network whose physical cost function has the form of Eq. 12, where *x*_*a*_ is the physical degree of freedom for node *a* and the learning degrees of freedom *w*_*i*_ are organized in the Hessian *H*_*ab*_ = *W*_*ab*_. These learning degrees of freedom evolve by Eq. 13. (2) A flow or resistor network where the physical degrees of freedom are the node pressures or voltages and the learning degrees are the edge conductances. The physical cost function is described by Eq. 14 and the learning rule is described by Eq. 16. (3) A mechanical central-force spring network, where the physical degrees of freedom *x*_*a*_ are the node positions and the learning degrees of freedom are the spring constants of the edges. Again, the physical cost function is Eq. 14 and the learning rule is Eq. 16.

## III. TRAINED NETWORKS ARE PHYSICALLY MODIFIED BY LEARNING

Above, we demonstrated a method for training physical networks to perform arbitrary functions in response to small inputs. We will show next that training modifies the physical properties of the network, changing the effective conductances, the dimension of the physical responses, and the alignment of the inherent coordinates of the physical response and the learned behaviors.

### A. Changes to the Hessian and its eigenspace

Above we discussed how to change the learning degrees of freedom to minimize a learning cost function in the response of a physical system. Suppose the system implements these learning dynamics. Then the changes to its responses to external forces are all manifested in modifications to the Hessian at the native state and its eigenspace. The Hessian changes for two reasons: (a) it is a function of the network weights 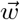 which are changing, and (b) it is evaluated at the native state 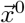 which minimizes the physical cost function of the network, and hence is implicitly a function of the changing weights 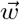. To evaluate these changes we first define the physical Hessian at a general network state 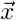 as 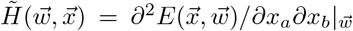. Evaluating 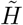 at the native state 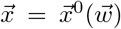, which minimizes the physical cost function, gives the Hessian 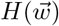 in (1).

In terms of 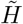 the change in the Hessian can now be written as

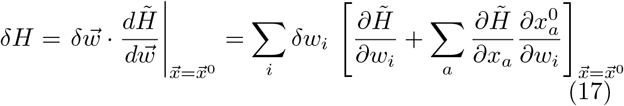

In the right hand expression the sum on *i* in the second term gives 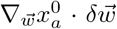, which is the change in the native state variable 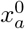 driven by learning. The expression 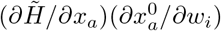 contains an implicit inner product in the physical space if the node variables *x*_*a*_ are regarded as d-dimensional vectors (see Sec. II B.)

If the physical cost function is quadratic in the node variables 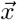 like in (12) or (14), or we are in the linearized regime of small forces with a good quadratic approximation to the physical cost function (1), then 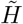 is by construction independent of 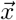, and 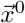 is independent of 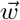. In these cases, which we will mostly study, the second term in (17) vanishes (or is small relative to the first term), and can be dropped. Such physical systems are in fact common: for example, electrical resistor networks, as in Ref. [27] always remain in the linear regime. Flow networks remain linear in the low Reynolds number regime, while initially unstrained spring networks are also linear over a reasonable range of strain when learning is performed on spring stiffness. In such systems the native state 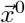 does not depend on the learning degrees of freedom at all, and stays fixed throughout the learning process.

As before, let us first discuss fully connected linear node networks – physical systems where the node degrees of freedom are trained to show desired responses. The change in the Hessian of such learning systems is given by Eq. 13:

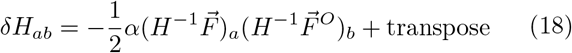

where we used the fact that the Hessian in this case equals the edge weight matrix *W*.

We see that the change in the components of the Hessian is given by a rank-1 symmetric matrix. This is the modification of the Hessian given a single task, i.e., input force output constraint pair. Each additional task that the network is trained for contributes such a modification, and therefore the total change in the Hessian due to a learning step consisting of *r* = 1 … *n*_*T*_ tasks is

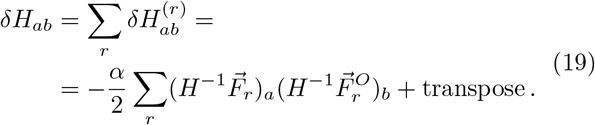

Recalling that 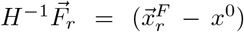 and 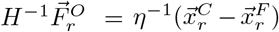, we see that Eq. 19 is a symmetrized outer product of the input response and the output nudge:

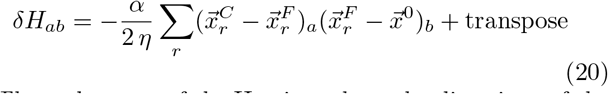

Thus, elements of the Hessian where the directions of the input and output responses agree tend to be reduced, and otherwise tend to increase. In particular, if the two responses are parallel (anti-parallel), the effect of learning will soften (stiffen) the Hessian. Note that *δH* is a symmetric matrix, preserving the symmetry of the Hessian. However, this learning rule does not guarantee that the Hessian remains positive semi-definite.

Now that we know how learning modifies the Hessian, we can track its evolution and predict the important features of the system in the neighborhood of the native state. To do so, we first discuss how the eigenspace of the Hessian changes in response to learning. Using first order perturbation theory, the changes in the eigenvalues *λ*_*n*_ and eigenmodes 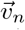 of the Hessian (*n* = 1 … *N*) due to one task (labeled *r*) in a learning step are:

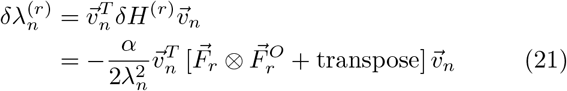

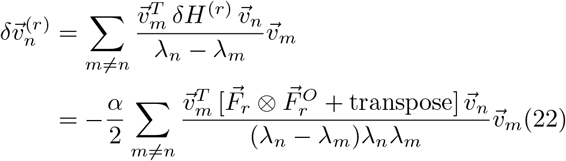

where the outer product ⊗ means that 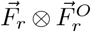 is a matrix with components 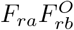, and we assume a generic starting network with non-degenerate eigenvalues. If the network is fine tuned to have some identical eigenvalues we will have to employ the techniques of degenerate perturbation theory, at least in the first learning step. The *λ*^2^ and *λ*_*n*_*λ*_*m*_ in the denominators of (21) and (22) arise from the action of the *H*^*−*1^ factors in *δH* (see Eq. 19) on the eigenvectors 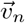 and 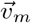 in the numerators. Note that the second order correction to the eigenvalues is 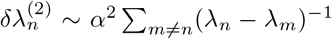 [51], and is thus neglible compared to the first order term (∼*α*) in the limit of slow learning rate *α* ≪ 1, except when two eigenvalues nearly cross. At that point an effective “repulsion” prevents eigenvalue crossing.

Two key points about the eigenvalue correction are: (1) Changes occur predominantly in the smaller eigenvalues because of the 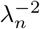 scaling, and (2) A fully trained system no longer changes its eigenvalues because, while the input force 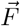 remains constant over training, the output force 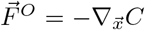 diminishes and vanishes together with the learning cost function.

We can also understand the eigenvalue shift geometrically. The projections of the input and output forces on the eigenspace tells us how these forces align with the different eigenmodes 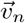. The contributions of the outer product 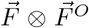 to the change in an eigenvalue are positive if both input and output forces align (or anti-align) with the associated eigenmode, and negative otherwise. Therefore, if the directions of the input and output forces both agree (or both disagree) with the direction of an eigenmode, its associated eigenvalue will decrease. Eigenvectors that lie (or have large projections in) the plane defined by the input and output forces are thus associated to strongly affected eigenvalues.

The correction to an eigenmode in Eq. 22 is a linear combination of the other eigenmodes. Note that eigenmodes corresponding to lower eigenvalues have a larger correction, and that close eigenvalues contribute larger corrections as well. For eigenmodes with the lowest eigenvalues the denominator of the sum in (22) will tend to be negative. Thus, the overall sign will be positive if the eigenvector has a positive projection onto both the input and output forces. Geometrically, this means that these eigenvectors will change by increasing their projection within the plane of the input and output forces. In other words, training effectively rotates the eigenspace so that the eigenmodes corresponding to low eigenvalues tend to align with the manifold spanning the input and output forces. This also suggests that the low eigenmodes rotate so that their associated eigenvalues are *effectively reduced* by the learning process, since (21) implies that eigenvectors aligned (or anti-aligned) with the forces reduce their eigenvalues during learning. Therefore, we expect that training the system for *any* task reduces the lower lying eigenvalues [36, 37].

For the upper eigenvalues of the Hessian the denominators of the sum in (22) tends to be positive, making the overall sign negative for eigenvectors with a positive projection onto the input and output forces. Thus, learning will cause these eigenvectors to rotate to align with either the input or output force, and anti-align with the other. In turn, this implies through (21) that the upper eigenvalues will tend to increase during training.

These considerations are verified in Fig. 3a, where we simulate the Hessian dynamics of fully connected linear networks with physical cost function (12) during learning of one task. We see that the bottom eigenvalue is significantly decreased, while the top eigenvalue increases in networks, here with *N* = 20 nodes, averaged over 50 realizations of networks and tasks. Moreover, by calculating the absolute value of the dot product between the eigenmodes and the input force 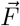, we find that the bottom and top eigenmodes align (or anti-align) with the input force, and hence with the trained task (Fig. 3b). Note that the eigenvectors are only defined up to sign anyway, so that alignment or anti-alignment is a matter of convention, and has no physical significance.

**FIG. 3.**
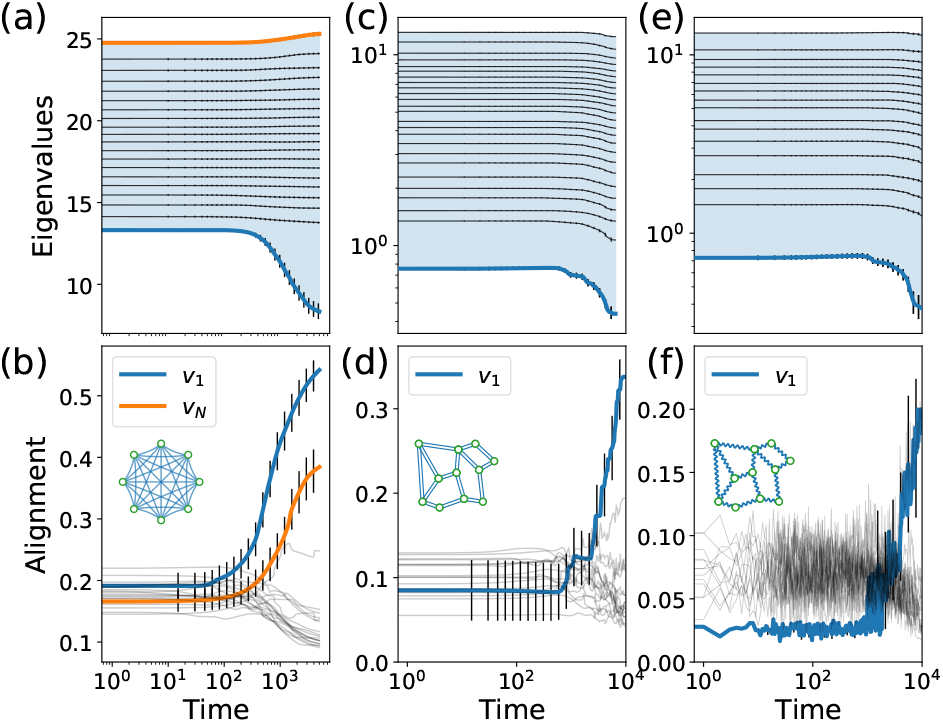
Hessian changes during learning. a) Hessian eigenvalue flow during training of fully connected linear networks (*N* = 20) for a single task. The lowest eigenvalue tends to be significantly reduced, showing that learning creates a soft mode. b) The lowest (blue) and highest (orange) eigenmodes of the Hessian significantly align with the input force defined by the task (measured by dot product). (c-f) Similar results are found for flow and mechanical networks (*N* = 40), except that the higher eigenmodes do not align with the task. Results averaged over 50 realization of networks and tasks.

In mechanical and flow/resistor networks, where input and output forces are more naturally trained in terms of differences between values at connected nodes, we have seen how the edge weights (conductances or stiffnesses) change in Eq. 16. In such difference networks, the Hessian is modified by a training step as

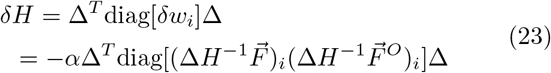

where we used the identification (15) between the weight matrix of node network and the edge weights of a difference network.

Here we wrote the change in the Hessian in terms of the product of responses of the network to the input and output forces. While more complicated than the result for node networks, this allows us to compute the change to the eigenvalues and eigenmodes of the Hessian similarly to Eqs. 21-22. Define the alignment matrix 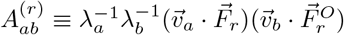 in terms of eigenvectors 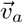 of the Hessian. This matrix captures the correlations in the alignments of the eigenvectors with the input and output forces, scaled by the eigenvalues. In terms of the aligment matrix, the change in the eigenvalues and eigenmodes of the Hessian, to first order in perturbation theory, is

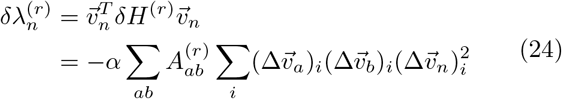

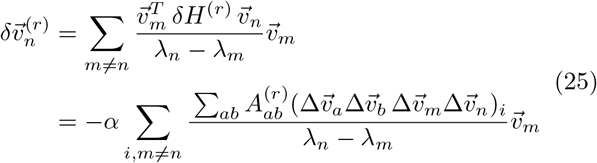

While this result is more complicated than the case of the node networks, the phenomenology associated with the low lying eigenmodes is similar. An eigenvalue *λ*_*n*_ is reduced if its associated eigenmode, after action by the difference operator, 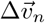, aligns with low lying eigenmodes that themselves align with the input and output forces. This causes the set of aligned eigenmodes with low lying eigenvalues to decrease their eigenvalues further. Similarly to node networks, the associated eigenmodes 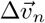 rotate to better align with the input and output forces. We conclude that physical difference networks such as mechanical and flow/resistor networks will respond to learning similarly to the node networks. One difference with the node networks is that all of the eigenvalues tend to decrease during learning. This is because the Δ operators in Eq. 24 causes all the shifts to carry contributions from the dominating low eigenvalues, which tend to decrease during learning. This is unlike the node networks where the alignment of each eigenmode with the input and output forces determines its own shift.

We verify these considerations for flow and mechanical networks with *N* = 40 nodes in Fig. 3c-f. The bottom eigenvalue is effectively reduced by learning and the associated eigenmode aligns significantly with the input force (results averaged over 50 realizations).

We note that machine learning algorithms do not typically have an analog for the physical Hessian, but are often concerned with a *cost Hessian* [52, 53] (the second derivative of the learning cost function with respect to the learning variables 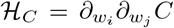). For trained learning models, this cost Hessian is typically low-rank, and has a number of stiff (non-zero) eigenvalues equal to the number of tasks the model was trained for. We discuss the cost Hessian ℋ_*C*_ and its relation to the physical Hessian *H* studied here in Appendix B, where we also show how the low-rank property of the cost Hessian arises in a physical learning setting.

Below we will discuss the consequences of the changes in the physical Hessian for physical properties of the learning system. For simplicity of notation, we will analyze node networks but the results will apply to flow/mechanical networks as well.

### B. Effective conductance

The properties of a physical system are often characterized by its responses to generic forces (e.g., finite temperature fluctuations), quantified below by an *effective inverse-stiffness/conductance* 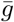. Suppose we compute the responses to *M* random forces 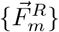 sampled from some distribution and indexed by *m*. In each case we have the free state response 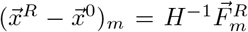. The effective conductance is the average amplitude of these responses:

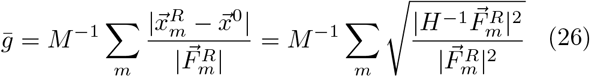

Now suppose that the random forces are drawn component-by-component independently from a Gaussian distribution and normalized to amplitude 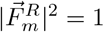. Decompose the Hessian as *H*^*−*1^ = *v*Λ^*−*1^*v*^*T*^ (*v* is a matrix of eigenmodes and Λ a diagonal matrix of eigenvalues *λ*). The eigenmodes *v* are a set of orthonormal vectors completely uncorrelated with the random forces. Thus the components of the vector 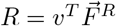 are inner products between random vectors on the unit sphere. In high dimension *N*, these inner products are approximately drawn from a normal distribution 𝒩(0, *N* ^*−*1^) with zero mean and variance 1*/N*. Now, note that

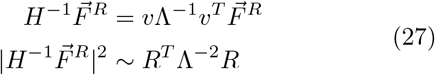

where in the second line we used the fact that *v*^*T*^ *v* is the identity. The second line in (27) equals the sum squared of the components of *R* scaled by the inverse square of the eigenvalues. Thus its expectation value is a scaled sum of the variances of the components of *R*, each of which equals 1*/N*. Putting everything together, we find that the expected value of 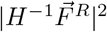 is a sum over the square inverse eigenvalues 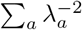.

Therefore, the effective conductance is a simple functional of the eigenvalue spectrum:

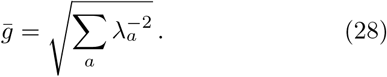

Note that the effective conductance 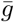 is dominated by the lower eigenvalues. It is expected to change during learning, as the eigenvalues *λ*_*a*_ of the Hessian change. In particular, we have seen that successful learning lowers the lowest eigenvalues, suggesting that trained systems have an increased effective conductance. Therefore, trained systems will be ‘softer’ than random systems, exhibiting larger responses on average to random applied forces. Note that the increased effective conductance is unrelated to the specific details of the learned task; such physical systems trained for *any* task are expected to become softer/more conductive.

Fig. 4a shows how the effective conductance of the different physical networks studied generally rises during training for a single task. These results are normalized such that the effective conductance at the beginning of training is 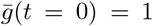, and averaged over 50 different realizations of the network and training task. Here, the orange curves correspond to flow networks, the green ones correspond to mechanical networks, and the blue curves correspond to the fully-connected networks.

**FIG. 4.**
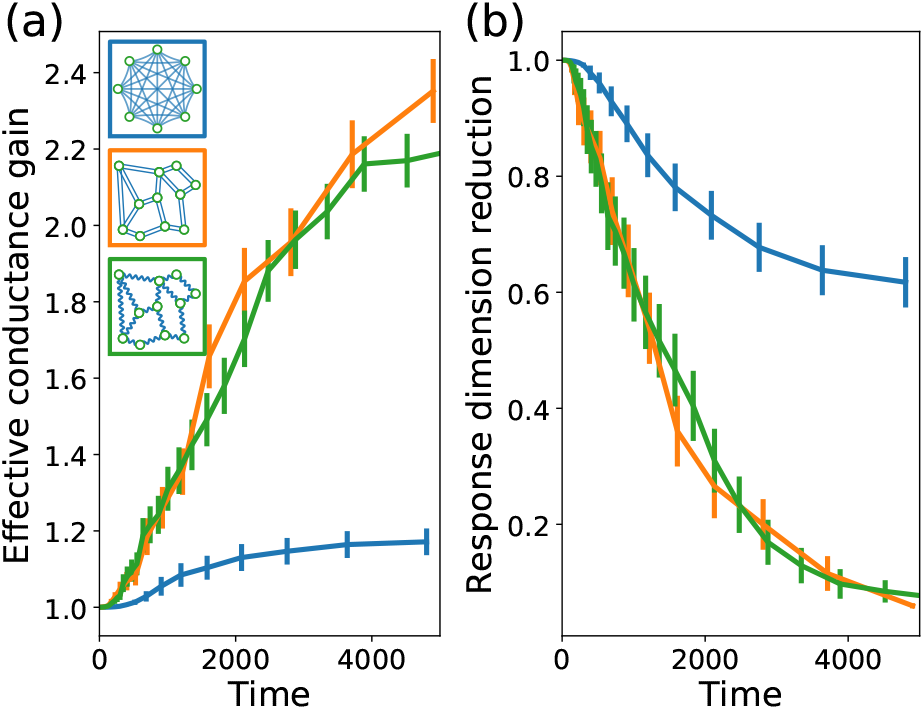
Training increases the effective conductance and reduces response dimension in physical networks. a) Effective conductance is increased during training in all systems considered, suggesting that trained systems are softer, with stronger responses to random forces. b) The physical response dimension is decreased in all systems during learning, so that the response of these systems to random forces is concentrated in low-dimensional manifolds. Results averaged over 50 networks and tasks.

### C. Physical response dimension

We can also directly study the dimension of the space of responses to random forces. While the system has *N* physical degrees of freedom, typical responses are coupled, lowering the effective dimensionality, as has been observed in, e.g., proteins [31] and neural circuits [46]. This effective dimension can be extracted by measuring how widely the responses are spread over the different eigenmodes of the Hessian.

To define a measure of this spread, we first consider the response 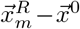 to a given random force 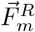. Let 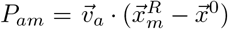 be the projection of the response onto the Hessian eigenmode 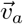. Then the associated *participation ratio* is

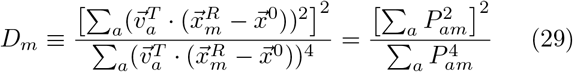

If a response 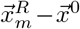 is parallel to a particular eigenmode 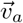, then it is orthogonal to the others, and so *D*_*m*_ = 1. On the other hand, if all eigenmodes participate in a given response with the same amplitude, i.e. 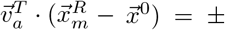 const for every mode, then *D*_*m*_ = *N*. Thus *D*_*m*_ captures the effective number of eigenmodes that participate in the response to 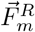.

In view of this, it is natural to define the effective dimension of the response as

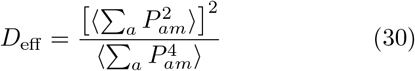

where the angle brackets denote an expectation value in the ensemble of random forces. In other words, we are defining the effective dimension as the ratio of the square of the second moment and the fourth moment of the projections of the response space onto the eigenmodes. Note that this is similar to, but not the same as, another classic measure of response spread: the participation ratio (PR) dimension [54–56]. The latter measure simply takes the expectation value of (29) in the ensemble of random forces, rather than separately taking expectation values in the numerator and denominator.

Our effective dimension has a simple and intuitive expression in terms of the spectrum of eigenvalues of the Hessian (details in Appendix C):

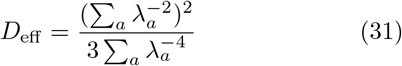

Notice first that if one eigenvalue is particularly small, it will dominate, and lead to an effective dimension of 1*/*3. As we showed above, learning changes the eigenvalues of the Hessian. To test how this changes the effective dimension we can take a derivative of (31):

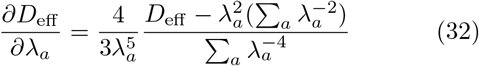

We see that changing a large eigenvalue does not modify the effective dimension much because of the 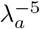 suppression. This makes sense because the system response to forces is controlled by the lowest lying eigenmodes – indeed (31) is dominated by the low eigenvalues. Conversely, decreasing the highest eigenvalues increases the effective dimension because this makes more eigenmodes active in the responses. In general, we saw that learning tends to decrease the lower eigenvalues of the Hessian eigenvalues, suggesting that learning reduces the physical response dimension for *any* learned task.

In fig. 4b, we compute the effective dimension during training for different physical systems (averaged over 50 networks and tasks). We find that the physical dimension generally decreases during training as the system adapts to accommodate the learned task, as expected given the Hessian eigenvalue dynamics seen before. There are other ways to estimate the dimension of the physical response – in Appendix C we discuss some of them and observe that for such alternative definitions, the physical dimension is still reduced by learning.

### D. Physical alignment with the learned task

As discussed above, the change in eigenmodes during learning tends to align the modes corresponding to low eigenvalues with the manifold defined by the input and output forces. The physical implication is that the system encodes information about the learned task in the eigenspace of its Hessian at the native state. In essence, the physical system, and particularly the native state basin, becomes the task it was trained for.

Remarkably, this allows an observer to glean information about a novel system without prior knowledge about whether this system was trained, and, if it was trained, for what task. Suppose we are given such a network that appears random to the naked eye. We can perform physical measurements, applying random forces to the network and measuring the effective conductance and physical response dimension described above. Comparing these values with those expected for a generic physical system of that type, we can tell whether the system was trained. But how can we deduce what task the system was trained for? The Hessian of the system can be measured in experiments by applying *N* orthogonal input forces, e.g., forcing each physical degree of freedom, and measuring the response, as 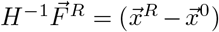. Then we can compute the eigenvalues and eigenmodes of the Hessian. Our analysis suggests that the lowest eigenmodes should have a large overlap with the response of the network to the input force, 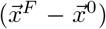, revealing the response function that the network was trained for.

For example, suppose that a network is trained to produce local responses at some output nodes in response to local forces at some other input nodes. Such a network is trained to strongly couple local inputs and local outputs. Therefore, we expect the eigenmodes related to this learned behavior to have large components in the physical degrees of freedom corresponding to the relevant input and output nodes. Measuring the Hessian and its eigenspace can thus inform us of where in the network these critical nodes are located, as well as what forces are expected at the inputs.

## IV. TRAINING FOR MULTIPLE TASKS

Above we saw how physical networks change when they learn a desired behavior in the small input force regime, where the physical response is described by the physical Hessian at the minimum of the native state basin. Learning changes the Hessian and its eigenspace to accommodate the learned task, aligning an eigenmode with the task induced coordinate system, lowering the associated eigenvalue, and creating a softer mode in the physical cost landscape. In this section we extend this reasoning to physical learning of multiple tasks in the same system, whereby the Hessian changes by averaging over single task modifications, as in Eq. 19. We thus expect the Hessian eigenspace to align with the coordinates of the different tasks. Since these tasks are in general independent from one another, training should result in aligning several Hessian eigenmodes (as many as the number of tasks). To verify this reasoning, we train fully connected node networks with *N* = 20 nodes and flow networks with *N* = 40 nodes to simultaneously satisfy five randomly sampled tasks (Fig 5, all results averaged over 300 realizations). In all cases, the tasks were learned well, resulting in vanishing error.

**FIG. 5.**
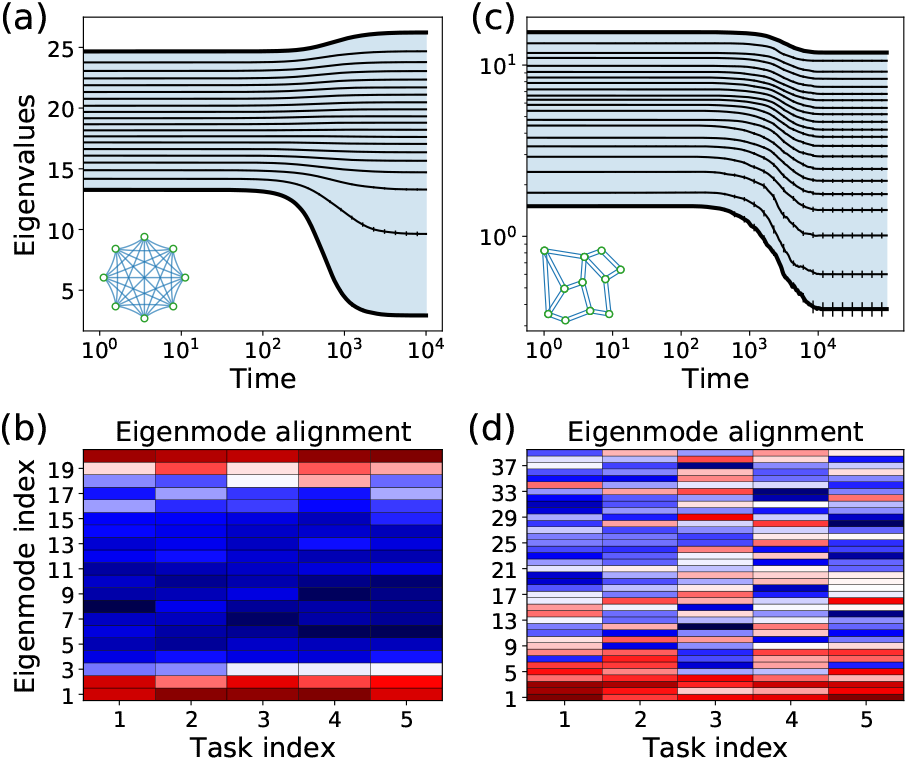
Hessian changes for systems learning several tasks. a) Hessian eigenvalues for a fully connected linear network trained for 5 independent tasks. A few of the bottom and top eigenvalues are modified significantly. b) Alignment (dot product) of the eigenvalues to the 5 tasks shows that ∼5 of the bottom and top eigenmodes align with the task inputs. c-d) Similar results observed when training flow networks for several tasks. For such difference networks the bottom eigenmodes tend to align with the input forces.

We see in Fig. 5a that several eigenmodes are significantly shifted in these networks. In the node networks, three eigenvalues are significantly reduced while two eigenvalues are increased. We also observe that five eigenmodes align (by dot product) with the five input forces (Fig. 5b), the bottom three and top two. In flow networks we see that all eigenvalues are decreased, with larger effects at the bottom of the spectrum (Fig. 5c). Furthermore, the bottom eigenmodes tend to align with the input forces (Fig. 5d).

It is, however, well-known that a system cannot be trained to perform too many tasks; physical and computational learning models have a *capacity M*_*C*_ for trained tasks. Trying to learn beyond capacity results in failure, where the system cannot successfully perform all of the desired tasks [7]. The capacity of simple learning models typically scales at best with the number of learning degrees of freedom (see Appendix D, where we argue our physical networks have a capacity that scales linearly with *N*_*w*_). This has been established for flow networks, where the number of output nodes that can be trained to respond to a single input is sublinear [49] in the total number of nodes, but can be raised to linear scaling [57] by avoiding frustration by tuning outputs in order of increasing distance from the source. We observe this finite capacity when training our models for multiple tasks (Fig 6a). Thus, we studied the physical effects of training beyond capacity.

**FIG. 6.**
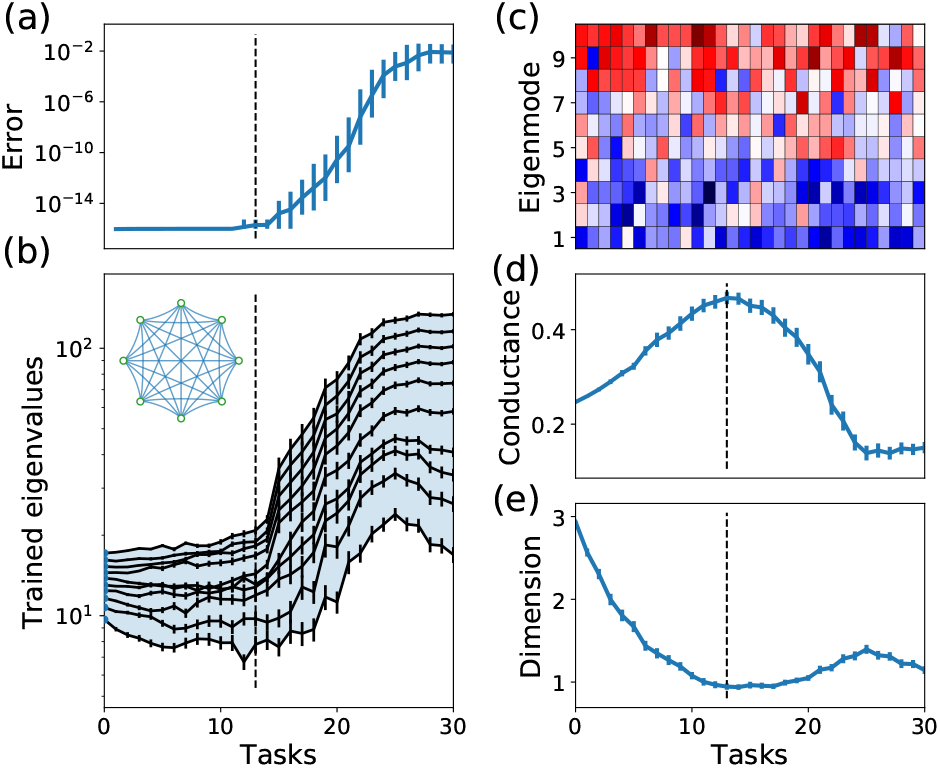
Training physical networks for multiple tasks. a) Training error remains low up until a capacity of tasks is reached. b) Eigenvalues of a network trained beyond capacity tend to increase. c) The bottom eigenvalues do not align with the tasks beyond capacity, while the top eigenvalues weakly align with them. d) Effective conductance as a function of the number of tasks system trained beyond capacity become less conductive (stiffer), as opposed to systems successfully trained below capacity. e) Physical response dimension remains low regardless of the number of tasks.

Consider the learning cost function for a task *r*, 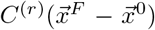. We defined the MSE cost function as a square of a constraint 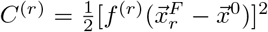. Since we are working in the small force regime, we can linearize any constraint in terms of the free state response as

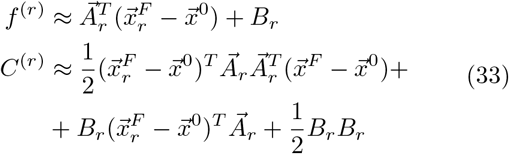

Next, noting that 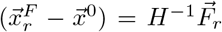, we can express the output force in the clamped state 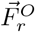 in terms of the input force 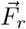:

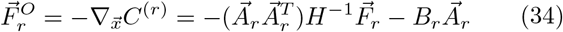

As discussed above, the modification of the Hessian and its eigenvalues due to learning involves the outer product

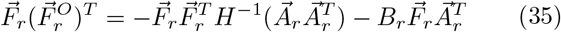

If we sum Eq. 35 over many random independent tasks 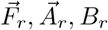, the second term in this sum averages to zero. However, as 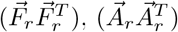 and *H*^*−*1^ are square, positive semi-definite matrices, the first term produces a nonvanishing contribution.

Now we observe from Eq. 21 that the change in the Hessian eigenvalues during learning is controlled by the quantity 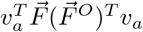 where we are projecting the outer product of the input and output forces onto the eigenvectors. To ask how the eigenvalues change for a typical set of tasks, we can average this quantity over an ensemble of random forces and task constraints. We take the components of the input forces and constraint elements to be independently sampled from zero mean, unit variance Gaussian distributions 𝒩(0, 1). Thus, the input and output forces are random relative to any given eigenmode *v*_*a*_. Then 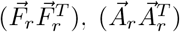 are random matrices, distributed such that their diagonal elements are drawn independently from independent chi-squared distributions 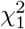, while their off-diagonal elements are drawn independently from 𝒩(0, 1). Thus, as 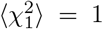, and ⟨𝒩(0, 1)⟩ = 0, these matrices average to the identity matrices. Since these two matrices are also independent, the average of *F* (*F*^*O*^)^*T*^ factorizes, and we have

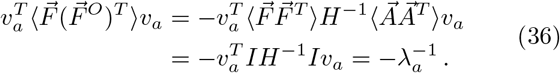

Using this result in Eq. 21, we see that the minus signs cancel and all eigenvalues are expected to receive positive adjustments that scale with the inverse eigenvalue 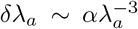. Thus, learning will make the system less conductive (stiffer). Below capacity, the Hessian eigenspace can align with the learned task(s), so that the lower eigenvalues can be effectively reduced. Above capacity, the Hessian cannot align with all the tasks, such that trying to learn some of them will tend to increase all eigenvalues. Eq. 36 gives the limiting behavior for averaging over an infinite number of tasks, but training over capacity is likely to cause an upward shift for most eigenvalues during learning.

This reasoning is supported in Fig. 6, where we train a fully connected node networks with *N* = 10 nodes for an increasing number of simultaneous tasks (all results averaged over 100 realizations of the network and tasks). In Fig. 6a, we plot the error after training as a function of the task number, demonstrating the finite capacity of the network (dashed line, *n*_*T*_ ∼ 13), defined here as the threshold above which some of the trained networks are unable to reduce the error to zero on all tasks. In other words we are defining capacity as the threshold in the number of tasks over which the learning process begins to fail. Fig. 6b shows the eigenvalues of the trained networks. As observed above, up to capacity the lowest eigenvalues tend to be decreased by learning. However, approaching capacity and beyond, the eigenvalues shift up during training, as suggested by Eq. 36. Examining the eigenmodes of systems trained beyond capacity at *n*_*T*_ = 30 (Fig. 6c), we see that the bottom eigenmodes no longer align with the trained tasks, but rather the top eigenmodes (weakly) align with them.

In Fig. 6d-e, we plot the effective conductance 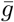 and the physical response dimension for systems trained for multiple tasks. The effective conductance is maximal (dimension minimal) when the number of tasks matches the capacity for simultaneous tasks as defined above (*n*_*T*_ ∼ 13). The conductance declines if we try to train additional tasks, and reaches a minimally conductive (stiffer) state when trained beyond capacity. In contrast, the effective dimension of the network responses remains low even beyond capacity. These results can be explained via Eqs. 28,31 by the observation that all eigenvalues of the Hessian tend to increase when training beyond capacity, not only the lower eigenvalues. The effective dimension is controlled by the *relative* size of the eigenvalues, and hence increasing all of them together does not change the dimension much.

Finally, we note that a physical system trained for multiple tasks can naturally be subject to noise during training: If the system learns in response to every observed task independently, the order in which tasks are presented can affect the learning process. Sampling training tasks randomly gives rise to a stochastic gradient descent-like algorithm [58], which can potentially affect the physical properties of the trained system as well. Moreover, physical learning is likely subject to other sources of noise, e.g. that affect the precision of the learning rules implemented by the system. In appendix E we study physical learning in such noisy conditions, and find that mild noise conditions, where learning is still possible, do not modify the physical effects discussed; the learning networks still find noisy solutions with high conductance and low physical dimension.

## V. DISCUSSION

In artificial neural networks, learning corresponds to traversing a path within the learning cost landscape that descends to a minimum. In physical learning systems, the physical cost landscape also changes as the system descends toward a minimum of the learning cost function, affecting the trajectory in the learning landscape. We chose to focus on the linear regime of small input signals where we can obtain the greatest insight into how learning restructures the physical landscape. We showed that networks can learn complex tasks ranging from digit recognition [21] to allostery already in this weak input regime. The structure of the response space is characterized by the spectrum of the physical Hessian around the minimum of the physical cost function, and the structure of the Hessian eigenmodes relative to the learned tasks.

While previous works demonstrated that training lowers low eigenvalues in the linear regime [33, 37], we have now traced the evolution of the eigenvalues and eigenvectors of the physical Hessian as learning emerges during the physical learning process. We find that physical learning necessarily leads to distinctive physical effects including strong responses to random inputs, as well as low-dimensional mechanical responses. This is remarkable as there are, in principle, many possible networks that satisfy a desired function without having this distinctive low-dimensional nature. If the number of tasks networks are trained for is below capacity (See Appendix D for more details on this capacity) they can essentially learn them all perfectly and the system can find multiple solutions. We showed that below capacity, networks physically evolve during training by becoming more conductive (less stiff in the case of elastic networks), lowering their effective response dimension, and aligning their eigenmodes with the learned tasks. In contrast, networks trained above capacity fail to learn and become *less* conductive (stiffer), but still maintain a relatively low effective response dimension. Thus an anomalously low network response dimension is a signature of learning, both below and above capacity. Our finding that training beyond capacity stiffens a physical system suggests a simple method of avoiding overtraining: as the trainer add tasks to a network, they should test its response dynamics to random forces, stopping when the stiffness begins to increase.

Our results suggest that low dimensionality is a generic outcome of physical learning in networks, possibly shedding light on the open question of why networks of neurons in the brain manifest surprisingly low dimensional response spaces. The generality of our approach further suggests a tool for analysing seemingly random physical networks to discover whether, and for what purpose, they are trained. Specifically, an experiment could measure the physical Hessian via small perturbations of the system, and then our results suggest that the eigenmodes corresponding to the lowest eigenvalues correlate to the tasks that the system was trained for. In other words, our results justify the naive intuition that the more responsive dimensions of a complex system encode its learned behaviors. Such tools can be useful in understanding newly discovered trained or evolved networks regardless of the specific details guiding their physical responses to perturbations.

In this paper we focused on the Hessian of the physical cost function around its minimum. In machine learning the focus is often on the Hessian of the *learning* cost or error function around its minimum (there is no analogy to the physical Hessian in most machine learning algorithms). In Appendix B, we show how the learning cost function Hessian, i.e., the second derivative of the learning cost function with respect to the learning degrees of freedom, also develops low-rank characteristics in a physical learning model. The Appendix also relates the learning Hessian to the physical Hessian, showing how physics encodes important information about the target functionality as a result of learning.

Our results were obtained in a scenario where the training data and the learning dynamics are noiseless. We tested that our findings are robust to the addition of Gaussian noise to the learning rule with a magnitude small compared to the learning rate (Appendix E). We also observe that introducing stochasticity in the selection of the order of training examples, or the order in which edges are modified, does not change our results. It would be interesting to extend our results to investigating noisier conditions when learning becomes challenging.

Finally, we note that we have shown that learning in the linear regime already has a remarkable phenomenology that can be analyzed powerfully. It would be very interesting to extend our results to nonlinear situations where the input and output forces are large. In this case, for example, the free state resulting from the application of an input may lie in a different basin of the physical cost function than the native state of the network in the absence of inputs. Other learning mechanisms can then come into play, such as the shifting of basins so that the minima themselves align with desired behaviors. In these cases, the learning rule may be uncorrelated with the native state, so that learning would not necessarily create soft modes lowering the effective response dimension [42]. However, the ubiquitous appearance of low dimensional response manifolds in systems that learn (see, e.g., [46]) suggests some of the findings might extend, perhaps in a different form, to physical learning with large inputs that explore multiple basins in the physical landscape.

## Acknowledgments

We thank Eric Shea-Brown, Giulio Biroli, Eric Rouviere and Sam Dillavou for insightful discussions. This research was supported by DOE Basic Energy Sciences through grant DE-SC0020963 (MS). VB and MS were also supported by NIH CRCNS grant 1R01MH125544-01 and NSF grant CISE 2212519. The Flatiron Institute is a division of the Simons Foundation, which provided support via Investigator Award # 327939 (AJL).

## Appendix A: Learning tasks (allostery, regression, classification)

In this work we simulate learning in physical networks with local, physically realizable learning rules, approximating the gradient of a learning cost function. The learning rules themselves are explored in the main text. Here we discuss some cost functions the networks can optimize, i.e., what tasks the networks can learn to perform. We describe the prototypical tasks used in the main text to explore the physical effects of learning. We then show that the physical effects are similar regardless of the type of task the network is trained to perform.

Each task explored for fully connected node networks in the main text is chosen as follows. An input force 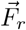 is randomly generated component by component from a zero mean, unit variance Gaussian distribution ∼ 𝒩(0, 1). Then an independent linear constraint 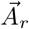 is also generated component by component from Gaussian distribution ∼ 𝒩(0, 1) along with a Gaussian distributed scale parameter *B*_*r*_ ∼ *𝒩*(0, 1). The linear constraint the network is trained to satisfy is then

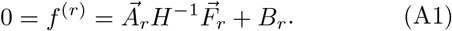

The associated cost function minimized by learning is 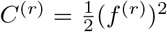. The results shown in the main text for fully connected node networks are based on optimizing such cost functions. In the linear response regime, these tasks form a basis for any desired functionality.

For difference networks, like the flow networks and mechanical networks studied here, these kind of tasks are readily learned – networks can often learn to satisfy such constraints with small modifications of the learning degrees of freedom, and hence small changes in the physical properties. To more clearly reveal the physical effects of learning, we challenged the difference networks to learn more difficult functions. The basic task we chose is *allostery*, which requires large target responses far away from source perturbations, a phenomenon previously observed in biological and mechanical networks [35] (see schematic in Fig. 7a). For each task, a random source node is chosen, and a force of magnitude *F*_*r*_ = 1 is applied as input at that node (in the positive direction for flow nets, and in some random direction in 2*D* for mechanical spring networks). Then, a random target node *o* is chosen, and the allosteric task is defined such that the response at that node has a finite amplitude 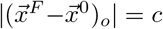 (*c* = 0.3 for flow networks, *c* = 0.5, and a random response direction is chosen for the 2*D* mechanical spring networks). When multiple simultaneous tasks are considered, we select multiple pairs of sources and targets, and apply such constraints to each pair. These allosteric tasks are more challenging for difference networks to learn, so that learning produces significant modifications to the network, and the physical effects discussed in the paper can be observed more readily.

**FIG. 7.**
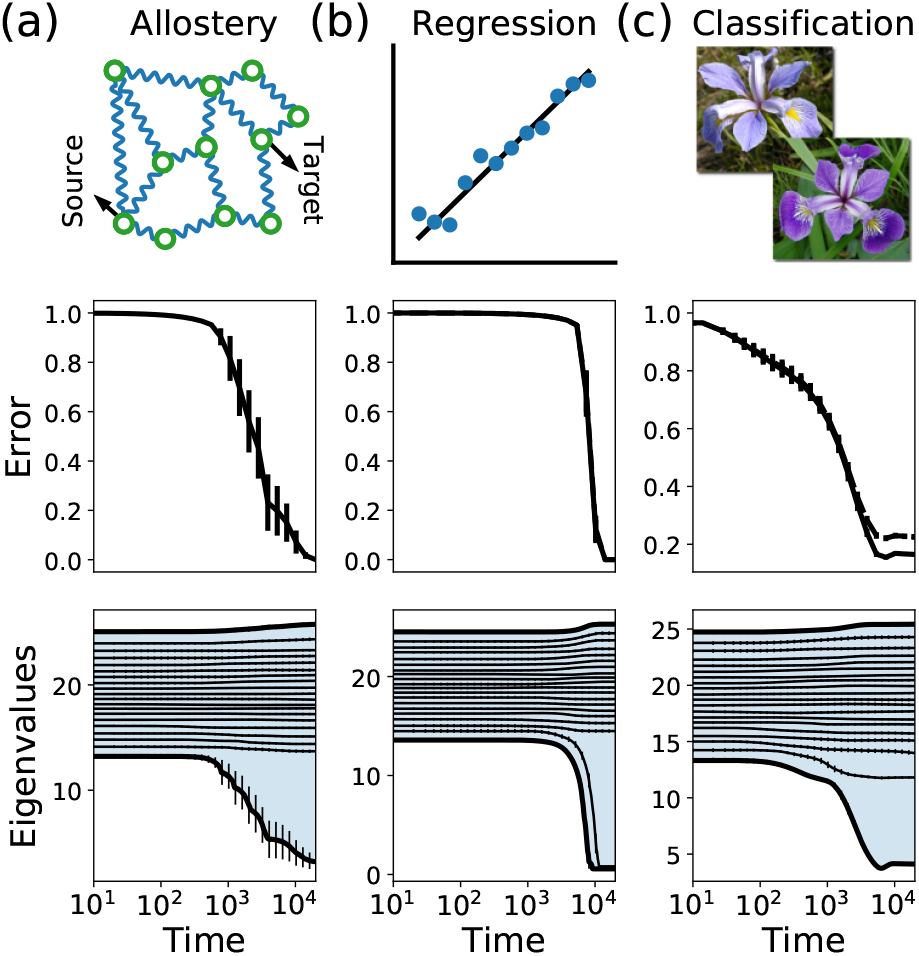
Training networks for different types of tasks. a) Allosteric tasks imply desired long range source-target relations. Here we show how networks can be trained for a single allosteric task involving one input and one output. Errors are shown in the middle row, and the dynamics of the Hessian eigenvalues on the bottom row. b) In regression tasks, the output units of a network are trained to recover a particular linear relation with the input units. Here we train networks for a set of two equations with two variables. c) Networks are trained to classify the Iris dataset [59] with 4 inputs and 3 classes (Iris species), using 10 out of 50 samples per flower for training. It does so by minimizing the cross entropy error, showing full line for the training error and dashed line for the test error. For all these tasks the physical learning phenomenology is similar; learning creates physical soft modes (results averaged over 10 networks).

To show that the physical effects of learning that we discussed are generic in the linear response regime, not only in terms of the physical network, but also in terms of the task(s), we train fully connected node networks for various types of tasks inspired by biology and computational machine learning. First, we train *N* = 20 networks on allosteric tasks similar to those defined for difference networks, with results shown in Fig. 7a (averaged over 100 realizations of networks and allosteric tasks). We find that training consistently reduces the error by orders of magnitude, and that the eigenvalues behave similarly to what we have seen earlier (Fig. 3a).

Other tasks of interest in supervised machine learning include regression, where the output of a network learns to recover some relation to the inputs based on training data. We train *N* = 20 fully connected linear networks to compute the results of a set of two linear equations. To do so, we randomly choose two input nodes *i*_1_, *i*_2_ and two output nodes *o*_1_, *o*_2_. Input forces correspond to the independent variables in the equations 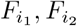, and the system is trained such that the response in the output nodes relates to the input forces as

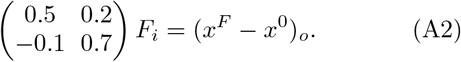

This is done by applying the learning rule of Eq. 10 for 50 random training forces, sampled from a Gaussian distribution as before. We further sample 100 *test* forces to measure the regression performance on data that is not used for training. Results are shown in Fig. 7b (averaged over 10 networks and training/test sets). The networks generally succeed in learning the desired input-output relation, decreasing the MSE cost by many orders of magnitude for both the training set (solid line) and test set (dashed line). More importantly, we track the eigenspace of the Hessian during learning, and find that two eigenvalues are strongly reduced when the network learns these two relations. Furthermore, the bottom two eigenmodes strongly align with the training set responses. This suggests the physical effects of learning are similar to the case of training for two tasks, discussed in the main text. Finally, we train networks for classification, where the system learns to assign labels to inputs. Networks with *N* = 20 nodes are trained to classify the Iris dataset [59], where the inputs correspond to 4 measurements of 3 species of Iris specimens. For each class of Iris we choose 10 specimens as a training set and the 40 others remain for a test set. The Iris measurements are applied as forces at four randomly chosen input nodes. A further three output nodes are chosen to correspond to the Iris classes. Since this is a discrete task (Iris specimens have discrete labels), the network ‘selects’ a label by having the largest response at the associated output node. Networks are trained by minimizing a cross entropy cost function that is more appropriate for discrete classification [24]. In essence, the modified learning rule is the same as Eq. 10 for specimens that are not classified correctly, while the output force vanishes for specimens that are classified correctly.

The results are shown in Fig. 7c (averaged over 10 realizations of networks and choices of training sets). As before, learning generally succeeds, significantly reducing the cross entropy error for both the training set (full line) and test set (dashed line). In terms of classification accuracy, the trained networks reach 100% training accuracy and 95% test accuracy. More importantly, the effects of learning on the physical network itself are again similar to the previous cases. The lowest Hessian eigenvalues are significantly lowered by learning, reducing the physical dimension and increasing the effective conductance of the system. We conclude that, in the linear response regime, the effects of learning on physical systems are generic and do not depend on the desired function.

## Appendix B: The cost Hessian

In this work we mostly studied the physical Hessian of a learning system and how it is modified by learning. Recently, the Hessian of the learning cost function 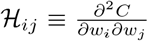 (i.e. the second derivative of the learning cost function with respect to weights *w*_*i*_) has received attention in the machine learning community. This Hessian was originally studied as a tool in improving gradient descent based techniques by including second derivative information [52], a procedure enabled by increased computing power. But studies have also shown that if a model is well-trained for *n*_*T*_ tasks, e.g., it has low training error for, say, *n*_*T*_ different classification classes, then the cost Hessian has *n*_*T*_ non-vanishing eigenvalues, while the rest of the eigenvalues vanish [53, 60].

Here we relate the physical and cost Hessians to see the origin of this phenomenon in physical systems. Note that the strong separation of scales in the cost Hessian is non-trivial, specifically as the physical Hessian has in general only soft modes and no zero modes (no vanishing eigenvalues). Furthermore, the physical Hessian *H* is a second derivative of the physical cost function with respect to physical variables 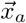, so the two Hessians live in different spaces and have different units.

Consider again the learning cost function for a network trained to exhibit *n*_*T*_ linear relations

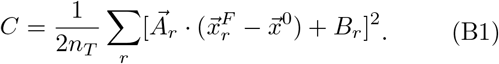

Let us compute the cost Hessian ℋ_*ij*_ of this cost function with respect to the learning degrees of freedom in several steps. First we compute the *Jacobian*, the change in the physical degrees of freedom (in the free state) given a change in the learning degrees of freedom:

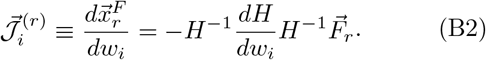

Here *H* is the physical Hessian and we used (3) for the free state response.

Choosing the fully connected node network as before, with some linear relation *H* = *Q*_*i*_*w*_*i*_, we have

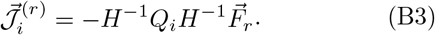

Next we calculate the second derivative of the free physical variables with respect to the learning degrees of freedom

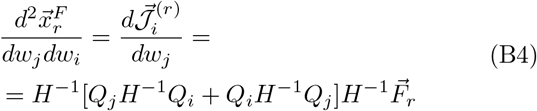

We are now in position to take derivatives of the learning cost function. Let us first find its gradient with respect to the learning degrees of freedom:

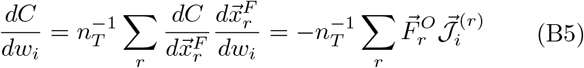

The cost Hessian is the second derivative:

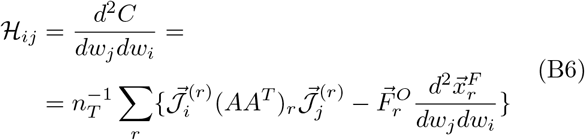

We can write down the cost Hessian in full form given our learning model, but Eq. B6 reveals its important features. Consider a well trained system, for which the value of the learning cost function tends to zero together with the output forces. In this case, the second term in Eq. B6 vanishes, as the system satisfies all the desired tasks:

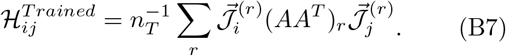

Note that Eq. B7 is a sum over tasks of outer products of the vectors 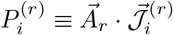:

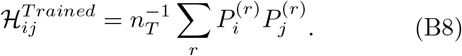

Since the matrices 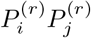 are outer products of a vector, we know that Rank 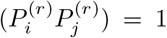. The total cost Hessian is thus a sum of *n*_*T*_ rank 1 matrices. By the sub-additivity of rank, it is guaranteed that the rank of the cost Hessian is at most *n*_*T*_. In other words, we expect the cost Hessian to have at most *n*_*T*_ non-vanishing eigenvalues, as observed numerically for machine learning and physical learning.

For completeness, we can also relate the physical Hessian and cost Hessian. Note first that:

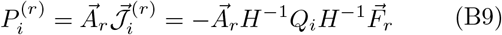

Thus, for well trained networks for which the training error vanishes, the cost Hessian relates to the physical Hessian as ℋ∼*P* ^2^∼ (*H*^*−*1^)^4^, suggesting that the low eigenvalues of the physical Hessian *H* dominate the *n*_*T*_ large eigenvalues of the cost Hessian ℋ.

## Appendix C: Physical response dimension

In this appendix we discuss measures for the physical response dimension, in particular the effective response dimension defined in the main text. Define a large set of *M*→ ∞ random forces 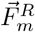, each of which sampled from the set of normalized vectors on the *N* -sphere, where *N* is the physical dimension of the system (number of physical degrees of freedom). The physical response of the system to these forces is given by

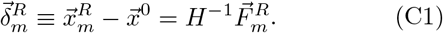

Applying the random forces, we obtain a set of *M* physical responses 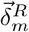. The Hessian of the native state can be decomposed to a set of non-negative eigenvalues *λ*_*a*_ and associated eigenmodes 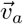 (whose number is equal to the number of physical degrees of freedom *N*). These eigenmodes correspond to different orthonormal ways in which the system can respond to perturbations. Thus, to estimate the response dimension, we can “count” the number of eigenmodes participating in the response.

Projecting the physical response over the Hessian eigenmodes gives the amplitude of each mode’s activation due to the external force 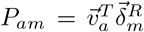. Summing this quantity over the eigenmodes and different random forces is self-averaging to zero because eigenmodes can be activated both positively and negatively. The physical response dimension is defined using these projections as an effective way to count the number of participating eigenmodes. For a given random force 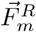 we have

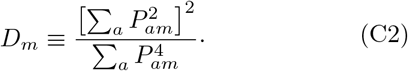

As discussed in the main text, this measure for the physical dimension has reasonable limits; it is *D*_*m*_ = 1 if only one mode participates, and *D*_*m*_ = *N* if all modes participate equally.

In view of this, we propose to characterize the effective response dimension by the quantity

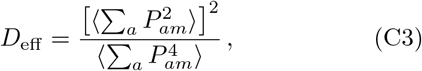

where the angle brackets indicate an expectation value over the ensemble of random forces. Thus *D*_eff_ measures the effective dimension as the ratio of the square of the second moment of the projections and the fourth moment of the projections.

We can compute *D*_eff_ by recalling that the inverse Hessian can be decomposed as *H*^*−*1^ = *v*Λ^*−*1^*v*^*T*^, where Λ^*−*1^ is a diagonal matrix of the inverse eigenvalues 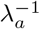, and *v* is a matrix who columns are the eigenvectors:

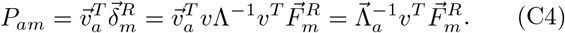

Here we used the fact that 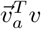 is a vector with zeros at all components, except a single 1 at component *a*. Therefore, only the *a*^th^ component of the vector 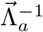 is nonzero, and equals 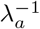. So, finally, we find that

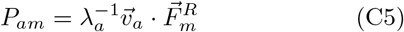

Next, let us consider the inner products 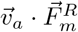. The set of eigenvectors 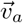 form a fixed orthonormal basis in *N* dimensions, while the forces 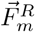 are random *N* dimensional unit vectors. Thus high dimensional systems (*N ≫* 1) the inner product of 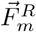 with any fixed 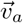 will be distributed according to a zero mean Gaussian with variance 1*/N*, 𝒩 (0, *N* ^*−*1^). Thus, over the ensemble of random forces, the second and fourth moments of *P*_*am*_ are given by

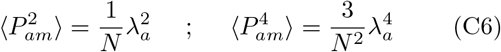

where we used the definition (C5) and the standard moments of a Gaussian distribution with variance 1*/N*.

Thus, the effective dimension in (C3) is

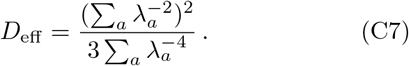

If one eigenvalue is very small and thus dominant, the effective dimension in response to random forces is 1*/*3 reflecting that fact that many forces will not drive the system much at all.

While in the main text we discussed *D*_eff_ as a measure of the physical response dimension of the system, there are several other measures of the intrinsic dimension of manifolds inspired by machine learning. We tested that our key qualitative results are independent of the choice of the measure of dimension. To do this we randomly select 500 normalized forces and applied them to systems during training (either fully connected linear networks or flow networks). We used the resulting responses to estimate the physical dimension using different methods: Manifold-adaptive dimension estimation [61], Correlation dimension [62], Maximum likelihood estimate [63] and the TwoNN algorithm [64]. Fig. 8 shows that the physical dimension is decreased during learning in physical systems regardless of the chosen method for dimension estimation.

**FIG. 8.**
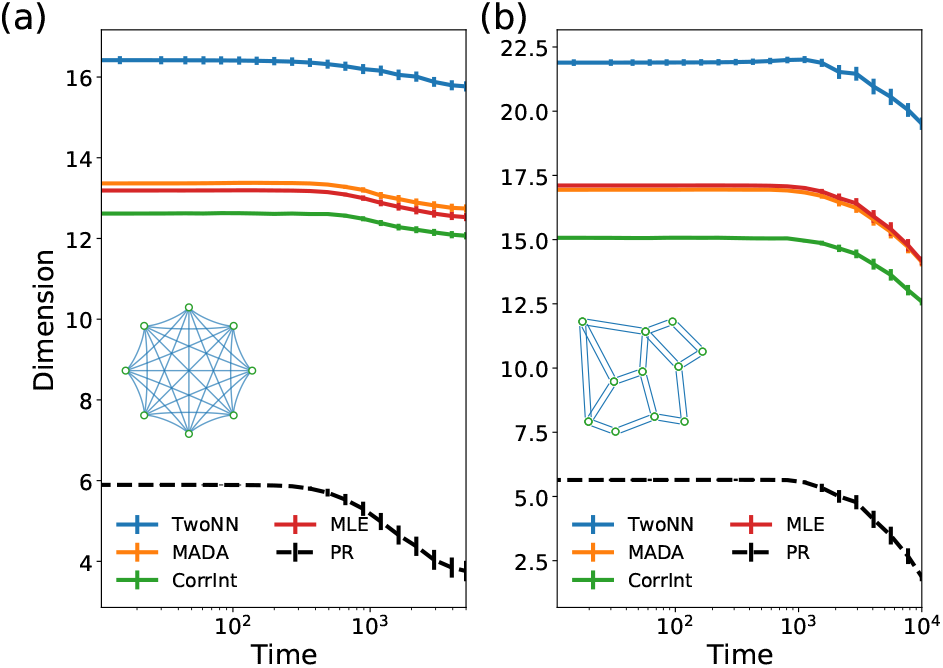
Different measures of the physical dimension. a) While in the main text we mainly used a measure derived from the participation ratio dimension to discuss dimensional reduction in the physical response, multiple other measures for the intrinsic dimension of the response manifold show the same result. Here we train fully connected linear networks and show how the physical dimension changes during training for several machine-learning-inspired measures. b) Similar results are found when training flow networks - physical dimension is reduced for all methods used to measure it.

## Appendix D: Learning capacity

In the main text we discussed how physical systems are able to learn multiple tasks up to a finite capacity. Here we argue that for systems trained in the linear response regime, this learning capacity is linear in the number of learning degrees of freedom *N*_*w*_, itself at most quadratic in the system size *N*.

For small input forces in the linear response regime, any learning cost function can be expressed as a sum of linear constraints, relating response of the physical degrees of freedom 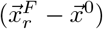 to an input force 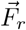:

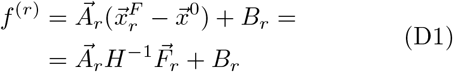

Successful learning means that *f* ^(*r*)^ = 0, or that for every task the system solves a linear equation

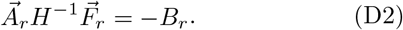

Note that the input force 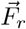, as well as 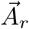, *B*_*r*_, are constants defined by the task to be learned, and the learning process only modifies element of the Hessian *H*. For a system with *N* physical degrees of freedom, we are allowed to modify the (at most) 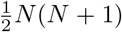 independent elements of the Hessian to find a solution to Eq. D2.

When training the system for *n*_*T*_ simultaneous tasks, each defined by its own set of input 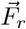 and constraints 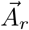, *B*_*r*_, the system has to find a solution to *n*_*T*_ independent linear equations at the same time. Learning attempts to modify the Hessian *H* such that all of these equations are satisfied simultaneously. As shown, the system has at most 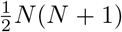 free parameters to satisfy *n*_*T*_ equations. A feasible solution exists only if the number of equations is at most equal to the number of free parameters. The network can thus learn a number of tasks which is at most equal to the number of independently trainable components of the Hessian. We thus expect a capacity proportional to the number of learning degrees of freedom, i.e., network weights, scaling at most quadratically with the system size and with a coefficient typically less than 1. This finite capacity is visible in Fig. 6a, where the error cannot be maintained at zero beyond *n*_*T*_ ≈ 13, which also marks a turning point in the effective conductance of the network.

Realistic physical networks are typically more constrained in the availability of adaptive (learning) degrees of freedom. In physical flow networks (e.g. vasculature) and mechanical networks (e.g. proteins), edges typically connect spatially neighboring nodes, so that the connectivity between nodes is short ranged, and the number of edges scales linearly with system size [16, 49, 57]. Thus their learning capacity is linear in *N*.

## Appendix E: Physical learning with noise

So far we discussed learning in pristine systems, where the learning rule is performed perfectly without any noise. In such cases, we found that physical learning can reduce errors arbitrarily (Fig. 2b) as long as the num-ber of tasks is below the network capacity. However, it is clear that real physical systems are prone to imperfections and noise, and that these issues can limit learning [27]. In this appendix we discuss the effects of noise on learning and its physical effects on learning systems.

A straightforward way to include imperfections in learning is to introduce additive Gaussian white noise in the learning rule (Eqs. 13, 16). We add such noise, with a small amplitude compared to the learning rate *α*, sampled from (0, (5 · 10^*−*3^*α*)^2^), and train fully connected node networks and mechanical networks for five independent tasks (Fig. 9). The noise in the learning rule produces a floor in the error achievable by the network (Fig. 9ab).

**FIG. 9.**
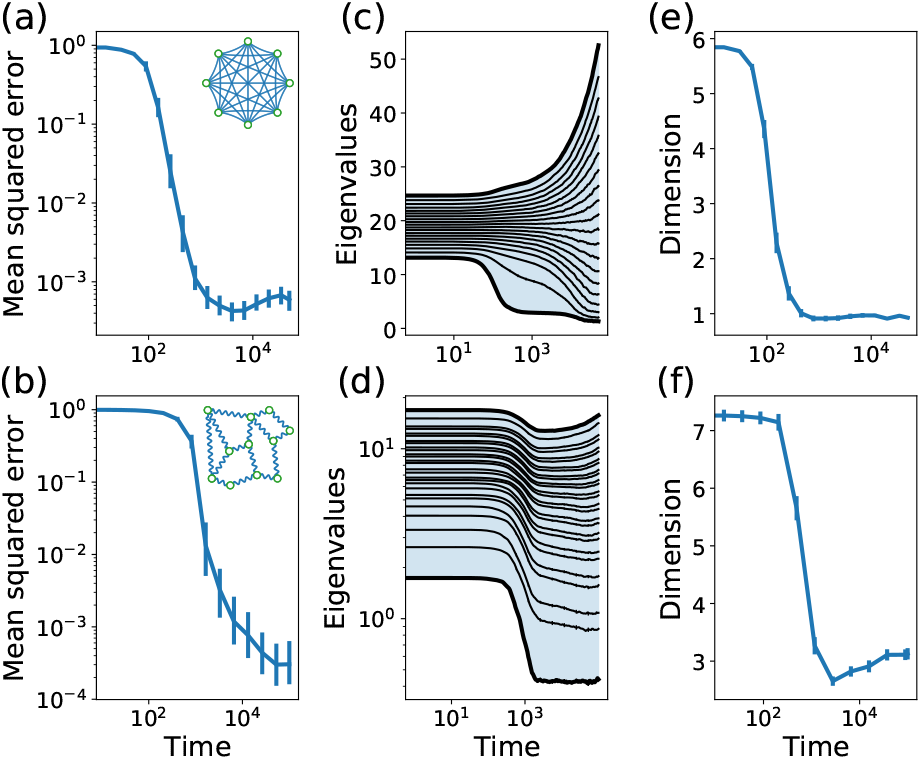
Noise limits the system’s learning ability, but the physical effects of learning persist. (a-b) Training fully connected linear networks and mechanical spring networks for 5 independent tasks with a noisy learning rule, we find that learning is still successful, yet is limited by an error floor. (cd) However, the dynamics of the eigenspectrum are weakly affected by the noise, dominated by a reduction of the lowest lying eigenvalues. (e-f) Therefore, the physical properties of the networks trained with noise remain similar to the noiseless case. In particular, we observe that training sharply decreases the physical response dimension of the network. All results are averaged over 100 sets of tasks.

However, this noise does not strongly affect the physical dynamics of the Hessian and its eigenspectrum; compare Fig. 9cd to Fig. 5. In this noisy case, learning still predominantly affects the lower eigenvalues, lowering them and aligning the associated eigenmodes with the task. We further see that the noise tends to raise the upper eigenvalues, but as shown above, these typically have only minor influence on the physical responses of the system. As the eigenspectrum dynamics are largely unaffected by this noise, we can verify that the physical effects of learning discussed previously also persist, in particular the physical response dimension (Fig. 9e,f).

Besides noise in the learning degrees of freedom that would exist in any physical or biological realization, different learning protocols may introduce additional sources of noise. For example, consider stochastic gradient descent (SGD), where the system is trained at each iteration for a subset of the tasks. Although SGD introduces effective noise in the training dynamics, it is known to perform implicit regularization and improve generalization in computational machine learning [58]. Another plausible source of noise is the fact that in biological learning systems, the learning degrees of freedom are not synchronized, each evolving independently from the rest [65]. We implement both SGD and the update desynchronization in our dynamics, letting the physical system train on one task and update 10% of the learning degrees of freedom at every learning iteration. While applying these protocols slows learning, we observe no qualitative difference in the physical properties of the trained system. We conclude that the physical effects of learning in the linear regime that we described in this work are robust to noise that will likely exist in experimental realizations, such as recent experiments in learning resistor networks [27].

